# Transcriptional trajectories of human kidney disease progression

**DOI:** 10.1101/342972

**Authors:** Pietro E Cippà, Bo Sun, Jing Liu, Liang Chen, Maarten Naesens, Andrew P McMahon

## Abstract

Our molecular understanding of clinical conditions progressing from acute organ injury to irreversible dysfunction is limited. We used renal transplantation as a model to characterize the transcriptional response along the transition from acute kidney injury to allograft fibrosis in humans. The integrated analysis of 163 transcriptomes with machine learning techniques identified shared and divergent transcriptional trajectories determining distinct clinical outcomes in a heterogeneous population. The molecular map of renal responses to injury was validated in a mouse ischemia-reperfusion injury model and highlighted early markers of disease progression. This generally applicable approach opens the way for an unbiased analysis of progressive diseases.

## Introduction

Genome-wide gene expression analysis has revolutionized biomedical research by providing a new level of understanding of human biology and by highlighting the molecular complexity of individual variability.^1^ The next step toward the goals of precision medicine is to decipher the molecular mechanisms determining the heterogeneity of multifactorial clinical conditions and to identify the best diagnostic and therapeutic targets to anticipate and prevent disease progression.^2^ Chronic inflammation and fibrosis are hallmarks of dysfunctional tissue repair leading to irreversible organ dysfunction, a process well exemplified by the progression of chronic kidney disease.^3^^-^^6^ Our previous studies in a mouse ischemia-reperfusion model demonstrated that the progression from acute kidney injury to fibrosis and irreversible damage follows a transcriptional program over months.^7^ The characterization of similar mechanisms in humans is hampered by the limited access to human tissue for time-course analyses and by the challenges related to the high variability of human biology and pathology.^1,8,9^ Despite additional elements of complexity specific to allograft transplantation, renal transplants offer a unique opportunity to study the response to tissue injury in humans. Every organ transplantation begins with ischemia-reperfusion injury in well-defined conditions at the time of surgery, and protocol biopsies – performed in many transplant centers – provide access to renal tissue over time. The recent advances in computational biology offer new opportunities to analyze large datasets. Machine learning techniques originally developed to analyze single cell gene expression are particularly effective in identifying similar transcriptomes in heterogeneous tissues and predicting transcriptional changes in intermediates in the transition between distinct biological states.^10^^-^^13^ The same computational approach might resolve the heterogeneity of clinical conditions with an unprecedented accuracy and facilitate the discovery of molecular mechanisms determining disease progression.

## Results

### Kidney allograft transcriptome variability across individuals and time

We evaluated the transcriptome of 163 protocol biopsy samples from 42 kidney transplant recipients at 4 time points: before implantation (PRE), shortly after the restoration of blood flow in the graft (POST), and 3 and 12 months post-transplantation. The clinical characteristics of the study population are presented in **Table 1**. The gene expression correlation analysis including the most variable genes in the whole dataset highlighted clusters of genes involved in a variety of biological processes from the immediate-early response after injury to organ fibrosis, with genes related to fibrosis and adaptive immunity forming a distinct cluster of highly correlated genes (**Figs. 1a-b**). Dimension reduction analysis by t-distributed stochastic neighbor embedding (t-SNE) differentiated an early (PRE and POST) and a late phase (3 and 12 months, **Fig. 1c**). Within the same phase, biopsies from the same patient tended to cluster together (**Suppl. Fig. 1**). The predominant role of inter-individual variability was confirmed by the gene expression variance decomposition analysis in linear mixed models: the variation in gene expression was greater among individual patients (21% of total variance in gene expression) than across time (9% of total variance) (**Fig. 1d**). Among the genes displaying a high level of variability among individuals, we confirmed previously reported genes with high inter-individual variability such as RPL9 and GSTM1.^1^ Thus, despite the inherent variability related to the clinical setting and to the analysis of small biopsy samples in the context of a biological process not necessarily homogenously distributed in the tissue, the substantial level of intra-individual transcriptional constancy in repetitive biopsies from the same kidney indicated that the transcriptomic profiles were reproducible, reflecting biological differences among patients.

**Fig. 1.**
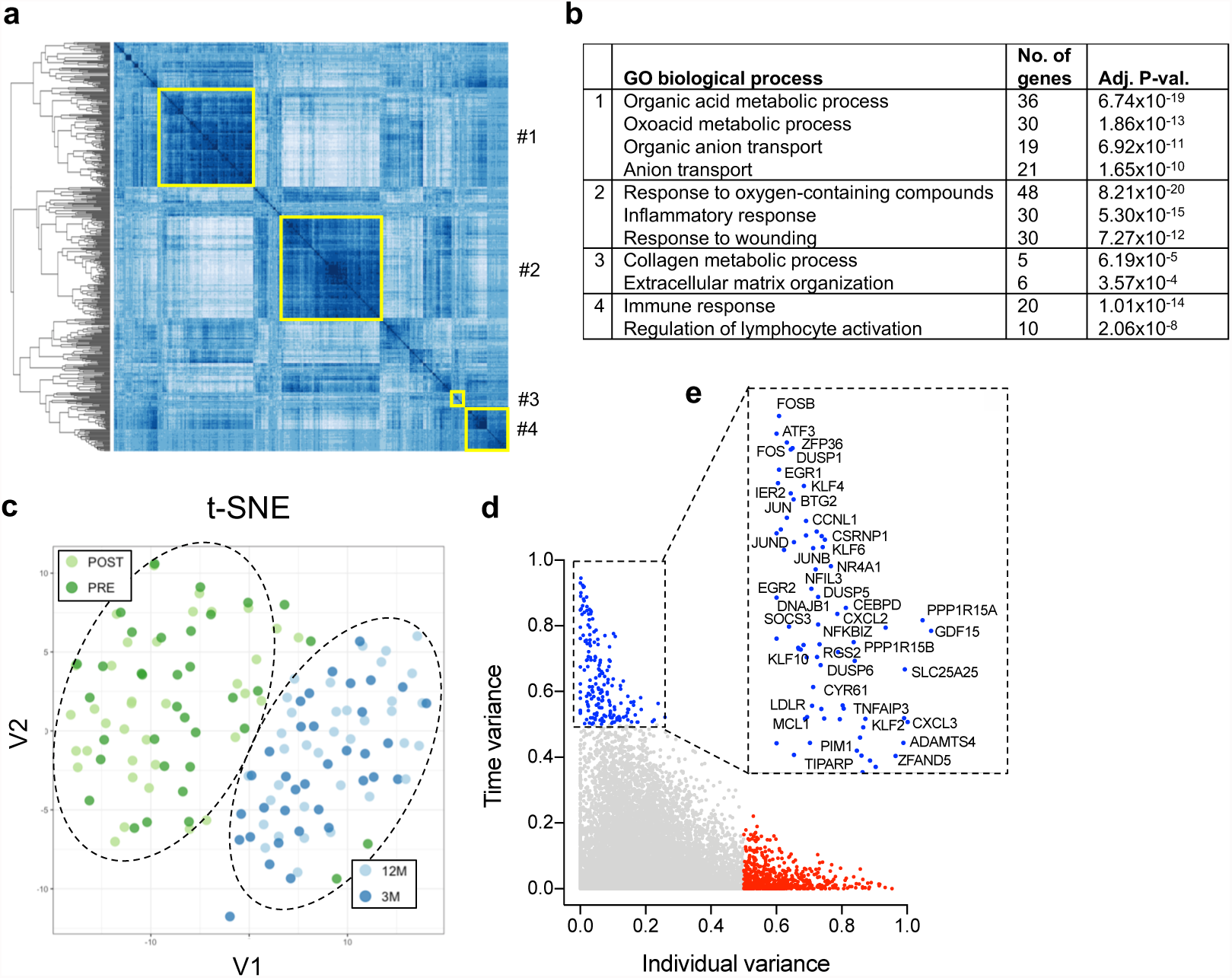
The kidney allograft transcriptome across individuals and time. (a-b) Gene expression correlation analysis including the 500 most variable genes across all samples;clusters of interest are highlighted in yellow and specified in (b). (c) t-distributed stochastic neighbor embedding (t-SNE) analysis on RNAseq data (RPKM values), including all samples and showing the separation of the transcriptomes in 2 major clusters:early phase (green), late phase (blue). (d-e) Contribution of individual and time to gene expression variation. Genes showing an individual-driven variance are shown in red, gene with a time-drive variance in blue and some relevant examples are specified in (e).

**Table 1.** Clinical characteristics of the study population. LD: living donor, DBD: donation after brain death, DCD: donation after cardiac death. Banff criteria: cg: chronic glomerular lesions, ci: chronic interstitial lesions, ct: chronic tubular lesions, cv: chronic vascular lesions. IFTA: interstitial fibrosis and tubular atrophy. ATN: acute tubular necrosis. TAC: tacrolimus, MMF: mycophenolate mofetil, CS: corticosteroids.

|  |  |
| --- | --- |
| Donor type (LD / DBD / DCD) (%) | 9 / 60 / 31 |
| Donor age (years) | 47.8 ± 14.9 |
| Donor sex (male / female / unknown) (%) | 45 / 53 / 2 |
| Donor creatinine (mg/dl) | 0.75 ± 0.24 |
| Recipient age (years) | 52.3 ± 13.0 |
| Recipient sex (male/female) | 64 / 36 |
| Recipient end stage renal disease diagnosis (%) |  |
| - Diabetes | 5 |
| - Hypertension | 14 |
| - Glomerulonephritis / Interstitial nephritis | 31 |
| - Congenital / Polycystic kidney disease | 21 |
| - Other | 12 |
| - Undefined | 17 |
| Previous renal transplant (%) | 7 |
| Cold ischemia time (h) | 12.2 ± 5.3 |
| Delayed graft function (%) | 14 |
| HLA-mismatches (mean) | 2.9 / 6 |
| Pre-transplant HLA sensitization | 14 |
| Histology of the kidney at baseline (PRE), according to Banff criteria |  |
| - cg (0 / 1 / >1) (%) | 100 / 0 / 0 |
| - ci (0 / 1 / >1) (%) | 63 / 29 / 8 |
| - ct (0 / 1 / >1) (%) | 34 / 66 / 0 |
| - cv (0 / 1 / >1) (%) | 82 / 11 / 3 |
| - ATN score (0 / 1 / >1) (%) | 3 / 19 / 16 |
| - % sclerotic glomeruli | 7.7 |
| Induction therapy |  |
| - Anti-CD25 therapy (%) | 36 |
| Maintenance immunosuppression |  |
| - TAC-MMF-CS (%) | 100 |

### Early transcriptional response to ischemia/reperfusion

The gene expression variance decomposition highlighted a small number of genes characterized by a low level of variability among patients but a high level of variability across time points (blue dots in **Fig. 1d-e**). This subset of genes was primarily involved in the immediate injury response (**Suppl. Fig. 2**), suggesting a low level of inter-individual heterogeneity in the initial response to ischemia (primarily triggered at the time point of tissue reperfusion).^14^ The reperfusion time, i.e. the time between restoration of blood flow and POST biopsy collection, was determined by the duration of the surgical procedure and was therefore variable among patients (range 26-88 minutes, **Suppl. Fig. 3a**). We considered this unsynchronized dataset as a time-course analysis of a biological transition, with each POST transcriptome representing an intermediate state along this process. Because of the analogy of this experimental setting with studies on cell state transitions investigated with single cell transcription analysis, we used a similar computational approach to order PRE and POST biopsies on a pseudotime trajectory (**Fig. 2a**).^12,15^ PRE and POST transcriptomes were correctly separated with PRE-samples classified in two groups (with differences in the expression of TIMP1 and in mitochondrial genes suggesting a different level of injury prior to reperfusion, **Suppl. Fig. 3b**) and all POST biopsies ordered along the same transcriptional trajectory (**Fig. 2a**). Consistent with the gene expression variance decomposition, the invariant sharp up-regulation of the early response genes in all POST samples (**Suppl. Fig. 3c**) and the absence of branching along the pseudotime axis suggested that all patients underwent a similar transcriptional program in the first hours after reperfusion. Interestingly, the order of patient biopsies along the pseudotime axis correlated well with actual reperfusion times documented in the patient’s clinical records (R^2^ = 0.69, **Suppl. Fig. 3d**). The integrated analysis of the transcriptional changes over pseudotime displayed a dynamic gene expression regulation: the first cluster of genes (**Fig. 2b**, cluster #1 in **Suppl. Fig. 3e, Suppl. Table 1**) was characterized by a rapid up-regulation and included immediate-early response genes (e.g. FOS, JUN and EGR1 in **Suppl. Fig. 3f**). This initial response was followed by a second wave of genes including the transcriptional regulators SOX9 and KLF5 (**Fig. 2b,** clusters #2-3 in **Suppl. Fig. 3e and g**).^16,17^ The regulation of other genes in the dataset can be interrogated online (https://lianglabusc.shinyapps.io/shinyapp). To identify the critical elements in the regulation of this coordinated process, we performed a network analysis, based on a modified Mogrify algorithm,^11^ which highlighted a major role for the transcription factors FOS in the context of a complex transcriptional network (**Fig. 2c, Suppl. fig. 4, Suppl. Table 2**).

**Fig. 2.**
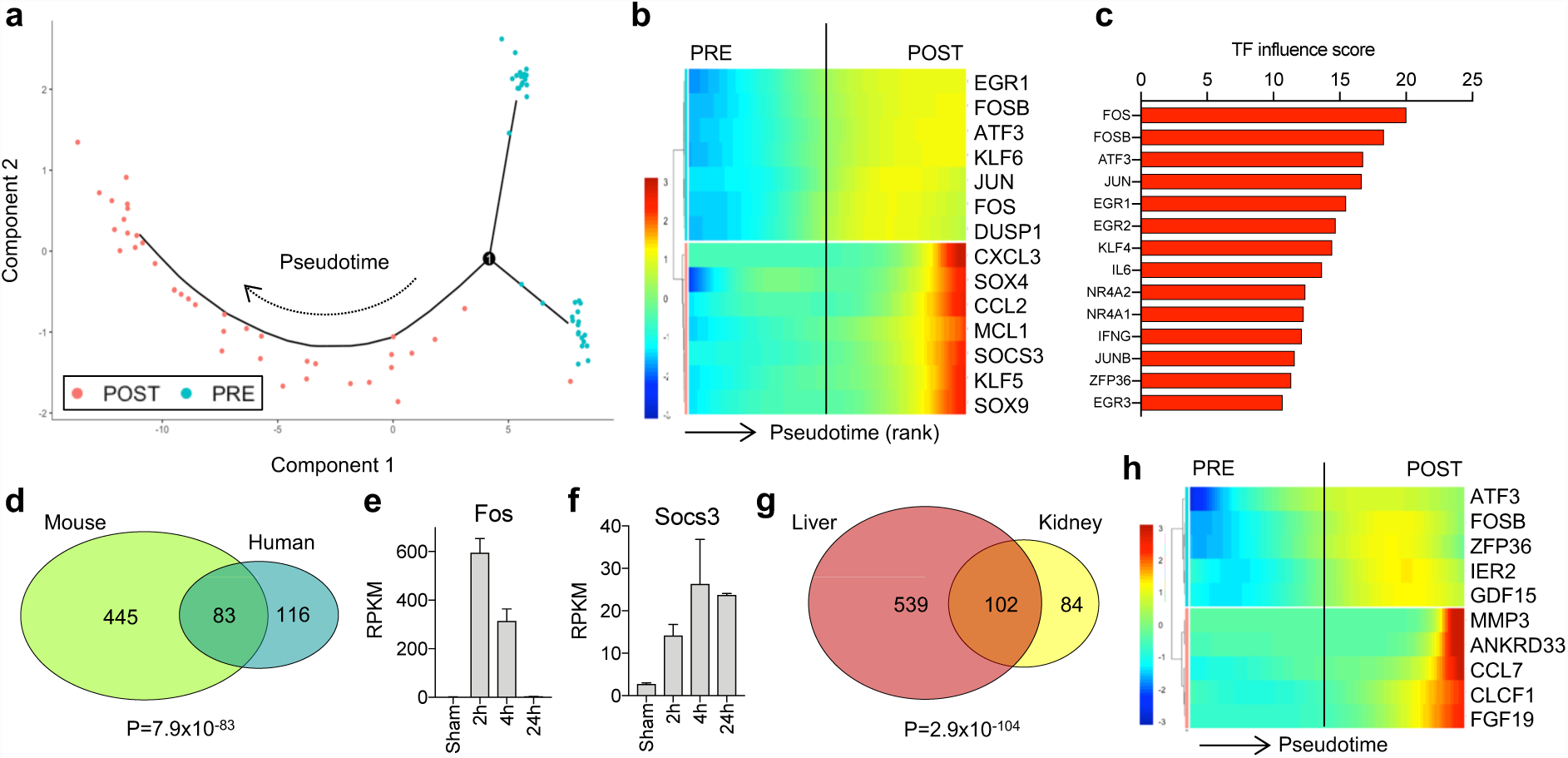
Early transcriptional response to ischemia/reperfusion. (a)Sample state ordering in the reduced dimensional space of PRE and POST samples, as determined by the Monocle algorithm. PRE samples were classified in two groups and are shown in cyan;POST samples ordered along a pseudotime line from right to left and are shown in red. (b) Cluster analysis of representative genes differentially expressed along the pseudotime:samples are aligned from left to right according to the order shown in (a). Genes are vertically aligned and classified in two clusters. The colors indicate the relative expression of the genes (log_10_ scale). (c) Influence score for the top 14 transcription factors as determined in the network analysis. (d) Venn diagram including genes differentially expressed in POST compared to PRE (human) and homologous mouse genes differentially expressed 2h after IRI compared to control. (e-f) RPKM values along the early time-course analysis after IRI in mice (N=3 for each time point). (g) Venn diagram including genes differentially expressed in POST compared to PRE in the human kidney and in the liver. (h-i) Cluster analysis of representative genes differentially expressed along the pseudotime in the liver.

The findings were validated in extensive data sets recently published by the group that characterized a mouse model of ischemia reperfusion injury.^7^ The comparison of differentially expressed genes in POST samples with homologous genes up-regulated 2h after IRI in the mouse model revealed a significant overlap across the species (**Fig. 2d**, **Suppl. Table 3**). Moreover, the time-course analysis in mice confirmed the same pattern of regulation of homologous genes. Genes classified in the first wave in humans reached the maximal expression level 2h after reperfusion in the mouse model (e.g. Fos, **Fig. 2e**), whereas the expression level of genes regulated in a second wave continued to rise at 4h (e.g. Socs3, **Fig. 2f**).

The general applicability of this approach was investigated by comparing the early transcriptional response to ischemia reperfusion across organs. We took advantage of RNA-seq data from liver transplants where liver biopsies were obtained before and after reperfusion.^18^ The comparison of the differentially expressed genes in PRE and POST in both organs highlighted a significant overlap (**Fig. 2g, Suppl. Table 5**). The liver dataset consisted of only 15 samples for each time point and the reperfusion time was less variable (ca. 2h). Nevertheless, pseudotime analysis delineated a biphasic transcriptional program (**Suppl. Fig. 5, Suppl. Table 5**): the first wave included the rapid upregulation of expression of mRNAs encoding transcriptional regulators governing the immediate-early responses. The overlap with the corresponding kidney dataset was less consistent in the second wave (12% vs 37%, P<0.0001), which was more organ-specific, identifying transcripts encoding signaling factors previously characterized as regulators of liver injury and regeneration (e.g. CCL7, FGF19)^19,20^ (**Fig. 2h**). Thus, as with previous experimental studies,^21^ the general pattern of transcriptional responses to ischemia-reperfusion was similar across organs, with a similar immediate-early response that likely initiates organ-specific mechanisms of injury and repair. More generally, an unsupervised computational approach generated a map of the early transcriptional response to ischemia/reperfusion injury in a real clinical condition reminiscent of regulated transcriptional networks previously investigated in cell culture.^22^

### Transcriptional trajectories of kidney disease progression

To identify groups of patients with a similar transcriptional profile at later stages post renal transplant, we first generated a minimum-spanning tree, based on global gene expression profiles at 3 and 12 months. The last 10 POST samples along the pseudotime presented in **Fig. 2a** were included as a control. We identified densely connected networks of samples (communities) by using random walks as implemented by the Walktrap community finding algorithm.^23^ Among the major communities identified with this approach, group A presented a transcriptional signature related to kidney injury, fibrosis and chronic inflammation (**Figs. 3a-b, Suppl. Table 6**). In analogy to the analytic approach applied for the early response, we considered the progression to fibrosis as a transitional process from acute injury (POST) to fibrosis (community A). The pseudotime analysis covering this transition correctly positioned the POST samples at one end of the pseudotime line (**Fig. 3c**). The late phase samples separated into two branches with all transcriptomes previously classified in the community A forming the latest stage of transition on the longer branch (**Fig. 3d**). In 86% of the patients both the 3 and 12 month biopsies were positioned on the same branch or moved to the branch depicting the progression to fibrosis over time, whereas only in 14% of the cases a transcriptome classified along the progression to fibrosis at 3 months was positioned on the opposite branch at 12 months. Early response genes displayed distinct characteristics in the long-term: a subset of genes (e.g. ATF3, IER2, FOSB) was restricted to the early response, whereas other genes showed a biphasic pattern, undergoing a secondary increase at late stages of chronic injury (e.g. SOX9, GDF15, **Suppl. Fig. 6**). Of particular interest was the late activation of HIF1A, consistent with an oxygen supply-demand mismatch in the fibrotic kidney,^24^ and JUN in consideration of its role in the pathogenesis of organ fibrosis (**Suppl. Fig. 6**).^25^

**Fig. 3.**
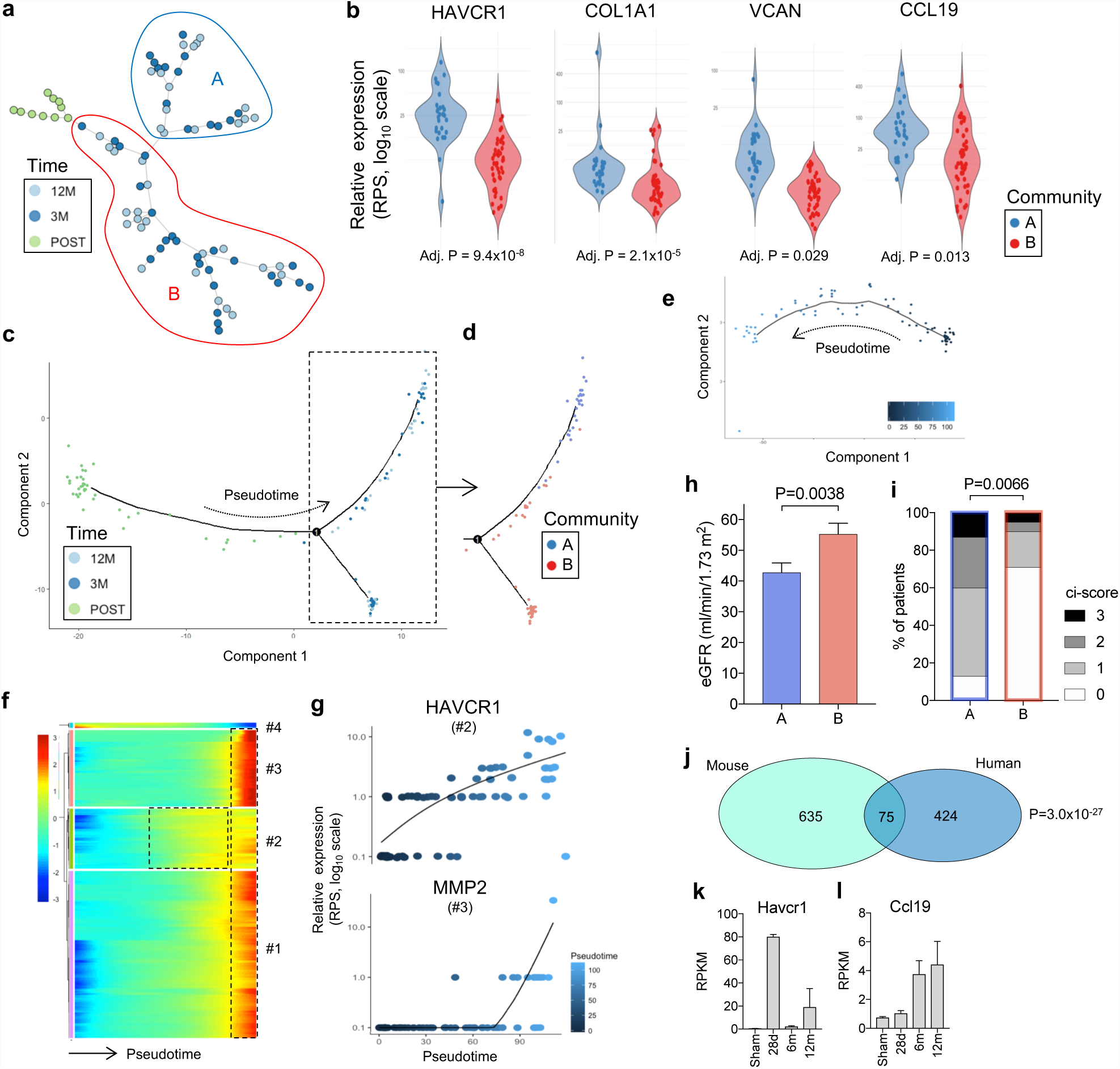
Transcriptional trajectory of transition from acute to chronic kidney injury. (a) Minimum spanning tree of kidney biopsy transcriptomes at 3 (dark blue) and 12 (light blue) months after transplantation, and 10 POST samples (green).Community A (marked in blue) separated from the rest of the study population (community B, in red). (b) Violin plots showing RPS values of representative genes selected as markers for kidney injury (HAVCR1, VCAN), fibrosis (COL1A1), chronic inflammation (CCL19). Adjusted P values are reported (Benjamini-Hochberg). (c-d) Sample state ordering in the reduced dimensional space of POST, 3 and 12 month samples, as determined by the Monocle algorithm. The colors indicate the time point of biopsy collection in (c) and the classification in community A or B in (d). (e) Similar analysis including only 3 and 12 month samples based on genes differentially expressed in community A and B. The color of the dots indicate the progression along the pseudotime, as indicated. (f) Cluster analysis of the top 2000 genes differentially expressed along the pseudotime shown in (e). Genes are vertically aligned and classified in 4 clusters (the complete list of genes is reported in the suppl. materials). (g) Representative example of one gene expressed early (HAVCR1) and late (MMP2) in the transition to fibrosis. The color indicate the pseudotime. The numbers indicate the cluster, according to (f). (h) Glomerular filtration rate at 12 months, estimated by CKD-EPI equation in community A and B. P value was calculated by Mann-Whitney test. (d) Histogram of the degree of fibrosis, quantified by ci-score on conventional histology. The % of patients in each fibrosis category in in community A and B is reported. The groups were compared by Chi-square test. (j) Venn diagram including genes differentially expressed in community A compared to B (human) and homologous mouse genes differentially expressed 12 months after IRI compared to control. (k-l) RPKM values along the late time-course analysis after IRI in mice (N=3 for each time point).

To specifically characterize the sequence of transcriptional events during the transition to chronic kidney injury we performed a pseudotime analysis including only samples collected at 3 and 12 months and based on genes differentially expressed in community A and B (**Fig. 3e**). Among the most significantly differentially expressed genes we found genes involved in cell death (e.g. DAD1, ANXA5) and complement regulation (e.g. CD59, SERPING1), which progressively increased with pseudotime.^26^ The upregulation of genes related to fibrosis (e.g. COL1A2, DCN, MMP2; clusters #1 and #3 in **Fig. 3f-g**) was a late event, coincident with the activation of genes regulating lymphocyte trafficking (e.g. CCL19, CCL21). Genes involved in the response to acute kidney injury (e.g. HAVCR1, LCN2) and innate immunity (e.g. TRAF6, TLR3) were upregulated earlier (cluster #2 in **Fig. 3f-g**). The cluster of genes down-regulated with pseudotime included mostly mitochondrial genes (cluster #4 in **Fig. 3f**, further characterized below). The complete list of genes and their distribution in each cluster is presented in **Suppl. Tables S7-8**. All differentially expressed genes along this pseudotime can be interrogated online (https://lianglabusc.shinyapps.io/shinyapp).

The functional relevance of this computational model was confirmed by reduced renal function, as determined by eGFR, and higher levels of fibrosis, as quantified by the ci-score on conventional histology, observed one year after transplantation in community A patients (**Fig. 3h-i**). Furthermore, in analogy to the interspecies validation of the early response, we compared genes differentially expressed in community A and B in patients with homologous mouse genes upregulated 12 months after ischemia reperfusion injury relative to sham-operated controls (**Fig. 3j**). Among the conserved genes across species in the late phase response, we identified genes associated with fibrosis (e.g. COL1A1, MMP2), adaptive immunity (e.g. CD2, IL2RG) and vascular biology (e.g. VCAM1, VWF) (**Suppl. Table 9**). Human-specific genes were mostly related to fundamental cellular functions indicating an additional human specific complexity in the response, whereas we did not find evidence for a specific regulation of genes related to adaptive immunity in humans, suggesting that the transplant specific elements exclusively present in humans were not predominant in our model. The sequence of events in the late phase in the mouse was similar as determined by the pseudotime analysis in humans, as exemplified by the intermediate expression of Havcr1 and the late up-regulation of Ccl19 (**Fig. 3k-j**). Thus, our analysis identified transcriptional trajectories describing the dynamic biological processes associated with the progression to chronic allograft dysfunction.

### Early discrimination of kidney disease progression

To identify early factors determining the alternative fate towards repair or progression to fibrosis we compared the transcriptomes classified right after the branching point (groups 1 and 2 in **Fig. 4a**). The first major difference among the groups was related to mitochondrial homeostasis: the expression of mitochondrial genes in group 1 was similar to normal renal tissue (defined by PRE samples obtained from living donors), but was significantly lower in group 2 and decreased further in group 3 (**Fig. 4b**). Additional evidence for a critical role of mitochondrial dysfunction and impaired ATP production in determining the discrimination between the 2 trajectories was provided by the analysis of key regulators of mitochondrial homeostasis (e.g. NRF1, PPARG, DNM1L) and glycolysis (e.g. PARP1) in the kidney (**Fig. 4c**).^27,28^ Conversely, the expression level of genes associated with innate immunity and extra-cellular matrix organization was higher in group 2, with the key regulator of kidney fibrosis MMP7 showing the strongest upregulation (**Fig. 4d, Suppl. Table 10)**.^29^ Among the genes displaying the highest influence score in the network analysis based on the comparison between group 1 and 2 we found TP53 and EP300 (**Fig. 4e, Suppl. Table 11**). EP300 was upregulated very early according to the pseudotime analysis presented in Fig. 3e-f (**Fig. 4f**). In line with the network analysis, the 33 genes most highly correlated with EP300 were sufficient for a classification of the transcriptomes in group 1 or 2 with a sensitivity and a specificity >90% (P<0.0001; **Fig. 4g, Suppl. Table 12**). Interestingly, binding sites of EP300 were recently associated with renal function in epigenome-wide association studies.^30^

**Fig. 4.**
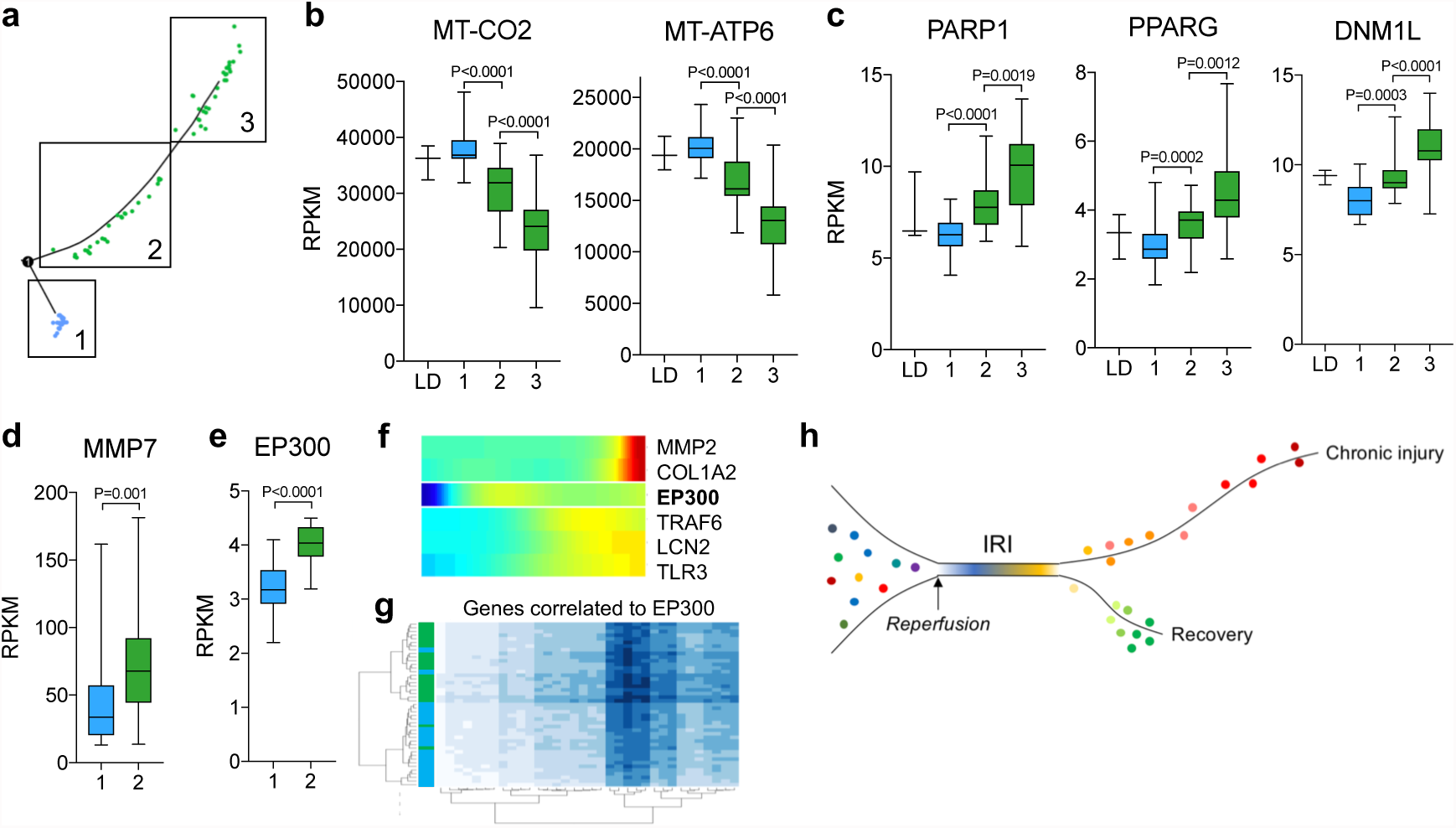
Early markers of transition to kidney fibrosis. (a) Schematic representation of the definition of groups along the divergent branches of the pseudotime analysis presented in Fig. 3e. (b-e) Box plots of RPKM values of the indicated genes in the group defined in (a), LD: living donor, as a surrogate of normal renal tissue. The groups were compared by Mann-Whitney test. (f) Expression profile of EP300 along the pseudotime presented in Fig. 3e-f in comparison with other genes. (g) Samples cluster analysis based on the 33 genes highly correlated with EP300 in groups 1 and 2 (Pearson correlation >0.88). The division of the samples in two clusters is indicated by the blue (group 1) and green (group 2) bars on the left. (h) Conceptual summary of the study highlighting the common early injury response after ischemia/reperfusion followed by a with divergent long-term outcomes:recovery versus the initiation of a chronic injury signature.

## Discussion

We investigated a complex clinical condition applying an unsupervised computational strategy, which integrates genome-wide expression analysis in heterogeneous groups of patients to identify and characterize shared trajectories of disease progression. Computational approaches originally developed to characterize *single cell* transcriptomes in heterogeneous tissues proved to be powerful tools to recognize and classify *single patient* transcriptomes in heterogeneous populations. The result is a molecular map of the transcriptional program following ischemia/reperfusion in the human kidney correlating with two major clinical outcomes: progression to chronic kidney injury or recovery (**Fig. 4h**). This approach has enabled a first-insight into the molecular mechanisms of individual human kidney disease progression. The early response to ischemia/reperfusion in the human kidney and liver confirmed in a clinical scenario identify similar waves of transcriptional regulation to those reported in time-course studies *in vitro.*^22^ Moreover, we identified early markers of disease progression, which represent potential targets to anticipate and prevent chronic kidney disease. The unsupervised computational analysis confirmed the clinical relevance of previous experimental studies, highlighting for example the critical role of mitochondrial dysregulation in the progression from acute to chronic kidney injury,^27^ and – at the same time - identified many genes of potential diagnostic and therapeutic relevance not yet investigated in this particular setting. The systematic analysis of gene-sets exhibiting similar behavior across species will facilitate the design of appropriately focused translational models in future mouse injury studies.^31^

Our data suggest the presence of common mechanisms in the response to ischemia/reperfusion and in the transition to chronic kidney damage despite the different factors contributing to potentially different types of kidney injury after transplantation in a heterogenous population. Thus, the molecular process characterized here reflects the fundamental response of the kidney to injury and the transition to organ fibrosis in humans. These fundamental processes might have a prognostic relevance and need to be considered for any further application of diagnostic strategies based on transcriptional analyses (including single cell transcriptomics). The overlap with the mouse IRI datasets indicates that many of these processes are conserved among species, and the overlap highlights those gene sets that are likely best suited for mouse-directed human disease modeling. Moreover, the presence of common elements in a non-transplant model expands the relevance of the human model beyond transplantation indicating that the molecular processes described here are mostly independent of the peculiarities related to transplantation (including unique immunological and pharmacological aspects).

Chronic, progressive diseases substantially contribute to global morbidity and mortality.^32^ Our computational approach can be applied to other organs and clinical conditions (as shown in the liver), and overcomes major limitations related to the need for repetitive tissue analyses over time to understand disease progression. In consideration of the level of accuracy obtained here even with relatively small numbers of samples, our approach might represent a powerful strategy to advance the molecular understanding of progressive clinical conditions.

## Acknowledgments

We thank Greg Alvarado, Jetty De Loor, Kari Koppitch, Gohar Seribekyan and Eric Tycksen for technical support. Work in APMs laboratory was supported by a grant from the California Institute for Regenerative Medicine (LA1-06536). PEC was supported by the Swiss National Science Foundation (grant 167773).

## Author contributions

PEC, MN, and APM contributed to conception and experimental design. JL performed IRI surgeries. PEC and JL performed experimental data acquisition and analysis. MN performed human data acquisition. PEC, BS, and LC performed human data analysis. PEC and APM prepared the manuscript, incorporating comments from other authors.

## Methods

### Study design, patients and clinical data

We performed a genome-wide gene expression profile by RNA sequencing (RNAseq) in kidney transplant recipients, randomly selected among patients with a full set of 4 available biopsies from a database at the University of Leuven. The patients received a kidney transplantation at the University of Leuven, Belgium. All patients gave written informed consent, and the study was approved by the Ethical Review Board of the University Hospitals of Leuven (S53364 and S59572). The renal biopsies were performed at the University Hospital of Leuven at following time points: before implantation (kidney flushed and stored on ice), after reperfusion (at the end of the surgical procedure), 3 months and 12 months after transplantation (protocol biopsies). Additional biopsies performed in the same patients for a clinical indication were not considered for this study.

For histological evaluation, kidney sections were stained with hematoxylin eosin (HE), Periodic Acid-Schiff (PAS) and silver methenamine (Jones). All biopsies were centrally scored by pathologists dedicated to transplant pathology following the same standard procedures. The severity of chronic histological lesions was semi-quantitatively scored according to the Banff categories for mesangial matrix expansion (“mm”), tubular atrophy (“ct”), vascular intimal thickening/arteriosclerosis (“cv”), interstitial fibrosis (“ci”), arteriolar hyalinosis (“ah”), and transplant glomerulopathy (“cg”). The team involved in the computational analysis was not informed about any clinical information until the end of the study, when computational and clinical data were matched. The histological evaluation was independent of the computational analysis and the pathologist was not informed about the results of the transcriptional analysis.

### Tissue storage and RNA extraction

Of each renal allograft biopsy included in this study, at least half a core was immediately stored on Allprotect Tissue Reagent^®^ (Qiagen Benelux BV, Venlo, The Netherlands), and after incubation at 4°C for at least 24 hours and maximum 72 hours, stored locally at −20°C, until shipment to the Laboratory of Nephrology of the KU Leuven. We performed RNA extraction using the Allprep DNA/RNA/miRNA Universal Kit^®^ (Qiagen Benelux BV, Venlo, The Netherlands) on a QIAcube instrument (Qiagen Benelux BV, Venlo, The Netherlands). The quantity (absorbance at 260nm) and purity (ratio of the absorbance at 230, 260 and 280nm) of the RNA isolated from the biopsies were measured using the NanoDrop ND-1000™ spectrophotometer (Thermo Scientific™, Life Technologies Europe BV, Ghent, Belgium). Before library preparation, RNA integrity was verified by high sensity RNA ScreenTape^®^ analysis. 5 samples were discarded because of not optimal RNA quality.

### Library preparation, RNA sequencing and data processing

Library preparation and RNA sequencing were performed in two batches. The first batch consisted of a pilot study with 3 patients (12 samples) and the second batch included the rest of the study population. To avoid biases related to batch effects, the samples included in the pilot study were excluded in the pseudotime analyses. The library was prepared with Clontech SMARTer technology at the Genome Technology Access Center of the Washington University, St. Louis, MO. The sequencing was performed in the same lab by using the HiSeq 3000 system on the Illumina platform, with a target of 30M reads per sample. The reads were aligned to the Ensembl top-level assembly with STAR version 2.0.4b. Gene counts were derived from the number of uniquely aligned unambiguous reads by Subread:featureCount version 1.4.5. Transcript counts were produced by Sailfish version 0.6.3. Sequencing performance was assessed for total number of aligned reads, total number of uniquely aligned reads, genes and transcripts detected, ribosomal fraction known junction saturation and read distribution over known gene models with RSeQC version 2.3.

### Data normalization and general statistical analysis

All gene expression levels were normalized and quantified by RPKM (number of reads per kilobase per million mapped read) and transformed into RPS (mRNA per sample) for pseudotime analyses (s. below). Dimension reduction analysis was performed with t-distributed stochastic neighbor embedding (t-SNE) by Rtsne package in R. Gene correlation analysis and feature correlation heatmaps were performed on PIVOT.^2^ Differential gene expression analyses were performed with EdgeR,^3^ by applying an exact test, with a false discovery rate of 0.05. P values were adjusted according to the Benjamini-Hochberg procedures. RPKM, RPS, eGFR values were compared by Mann-Whitney U tests. Categorical analyses were performed with Fisher’s exact or Chi-square tests. Venn diagrams were generated by the R package ‘VennDiagram’, and the significance of enrichment was calculated by hypergeometric tests. The community finding analysis was performed on PIVOT, as described by Darmanis et al.^4^

### Linear mixed model

To examine the gene expression variation that can be explained by the variables of interest a linear mixed model (LMM) was implemented with the R package lme4. Gene expression was modeled by variables of interest as random effects plus an intercept considered as fixed effects. In R syntax convention, the formula is:

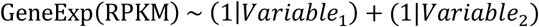

Variances were estimated by restricted maximum likelihood (REML) estimators. Then the variances explained by the two random effects were normalized by the total variance (sum of each variance explained by the random effect and the residual variance), which were used as the x-axis and y-axis coordinates. Genes with more than 50% variance explained by the variables of interest were colored. All 163 kidney samples, and only genes with maximum greater than 0.1 were kept with *time* (PRE, POST, 3 and 12 months) and *individual* as random effects. RPKM values were transformed as log10(RPKM + 0.01) before used in to the LMM model.

### Pseudotime analysis

All pseudo-time analyses were performed by using the Monocle workflow in R.^1^. First, genes with minimum RPKM greater than 0.1 and expressed in at least 40 samples were selected as *expressed genes*. RPKM values were then transformed into RPS by *Census* (included in Monocle 2).^1^ RPS was equivalent to RPC (mRNA per cell): We named it RPS since our data was bulk RNA-seq instead of single-cell RNA-seq. The minimum expression level for considering a gene in the Monocle cellset and differential expression analysis was 0.1, a negative binomial distribution was used. The differential expression analysis in Monocle was based on the time point of biopsy or on the classification in categories, as indicated in the text. Cell state ordering and gene expression pattern plots were generated with the codes included in the Monocle package.

### Network analysis

The network analysis was performed based on genes differentially expressed in PRE and POST samples. The analysis was performed in PIVOT by using the STRING protein-protein interaction database and the Regnetwork regulatory network database.^5^ The results were plotted with the igraph package.

### Comparison of mouse and human data

Mouse handling and husbandry and all surgical procedures were performed according to guidelines issued by the Institutional Animal Care and Use Committees (IACUC) at University of Southern California. The experimental model was previously characterized.^6^ Briefly, 10- to 12-week-old, 25- 28g, C57BL/6CN male mice, purchased from Charles River, were anesthetized with an intraperitoneal injection of a ketamine/xylazine (105 mg ketamine/kg; 10 mg xylazine/kg). Body temperature was maintained at 36.5-37°C throughout the procedure. The kidneys were exposed by a midline abdominal incision and both renal pedicles were clamped for 21 min using non-traumatic microaneurysm clips (Roboz Surgical Instrument Co.). Restoration of blood flow was monitored by the return of normal color after removal of the clamps. All the mice received intraperitoneal (i.p) 1 ml of normal saline at the end of the procedure. Sham-operated mice underwent the same procedure except for clamping of the pedicles. RNA was extracted from whole renal tissue with an RNeasy^®^ kit (Qiagen) and provided to the USC Epigenome Center’s Data Production Core Facility for library construction and sequencing. Library construction was carried out using the Illumina TruSeq RNA Sample Prep kit v2 through polyA selection. Libraries were applied to an Illumina flow cell at a concentration of 16 pM on a version 3 flow cell and run on the Illumina HiSeq 2000. Final file formatting, de-multiplexing and fastq generation were carried out using CASAVA v 1.8.2. The sequencing data were aligned to mm10 genome assembly with STAR aligner.

### Homologous reduction

The inter-species comparison was based on orthologous genes between human and mouse. The orthology information was downloaded from Biomart (https://www.ensembl.org/biomart) by selecting human data set hg38 and adding mouse orthology information. The Genome Assembly version used were GRCh38.p12 for human and GRCm38.6 for mouse.

### Comparison of kidney and liver

The liver dataset was downloaded from the GEO (https://www.ncbi.nlm.nih.gov/geo/query/acc.cgi?acc=GSE87487) and was analyzed in analogy to the kidney data.^7^

### Software

t-SNE, differential gene expression analysis, gene correlation analyses, the Monocle workflow, the community finding analysis and the network analysis were performed with the PIVOT platform, developed by the Kim Lab, University of Pennsylvania (http://kim.bio.upenn.edu/software/pivot.shtml). Violin plots were generated on the PIVOT platform and visualized on Plotly (https://plot.ly). Box plots and histograms were generated with Prism 7. Gene enrichment analyses were performed with ToppFun (https://toppgene.cchmc.org).

### Webpage

An interactive website was developed for a convenient interrogation of how genes of interest changed along pseudo-time in human kidney injury early response (PRE v.s. POST) and how genes behaved in the long run—a pseudo-time trajectory constructed from POST to 3 and 12 month samples.

---

**Suppl. fig. 1.**
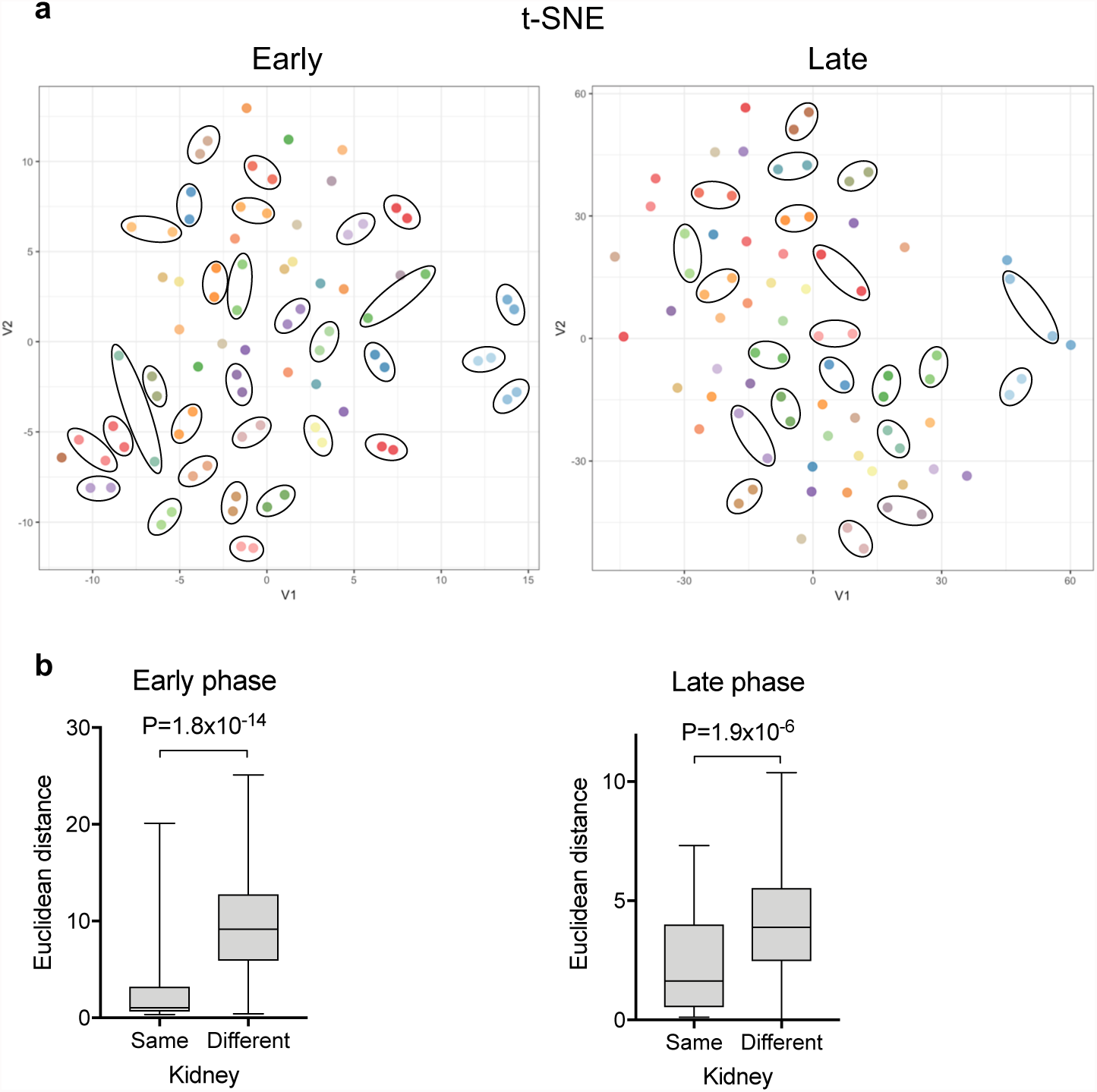
Intraindividual transcriptome variability. (a) t-distributed stochastic neighbor embedding (t-SNE) analysis on RNAseq data (RPKM values), including samples of the early (left panel; PRE, POST) or the late phase (right panel, 3M, 12M). Samples from the same patients are shown in the same color within each panel and are included in a circle if clustered together. (b) Box plots of the Euclidian distance between the points in the t-SNE analysis shown in (a) comparing the distance between the points from the same and from different kidneys, as indicated.

**Suppl. fig. 2.**
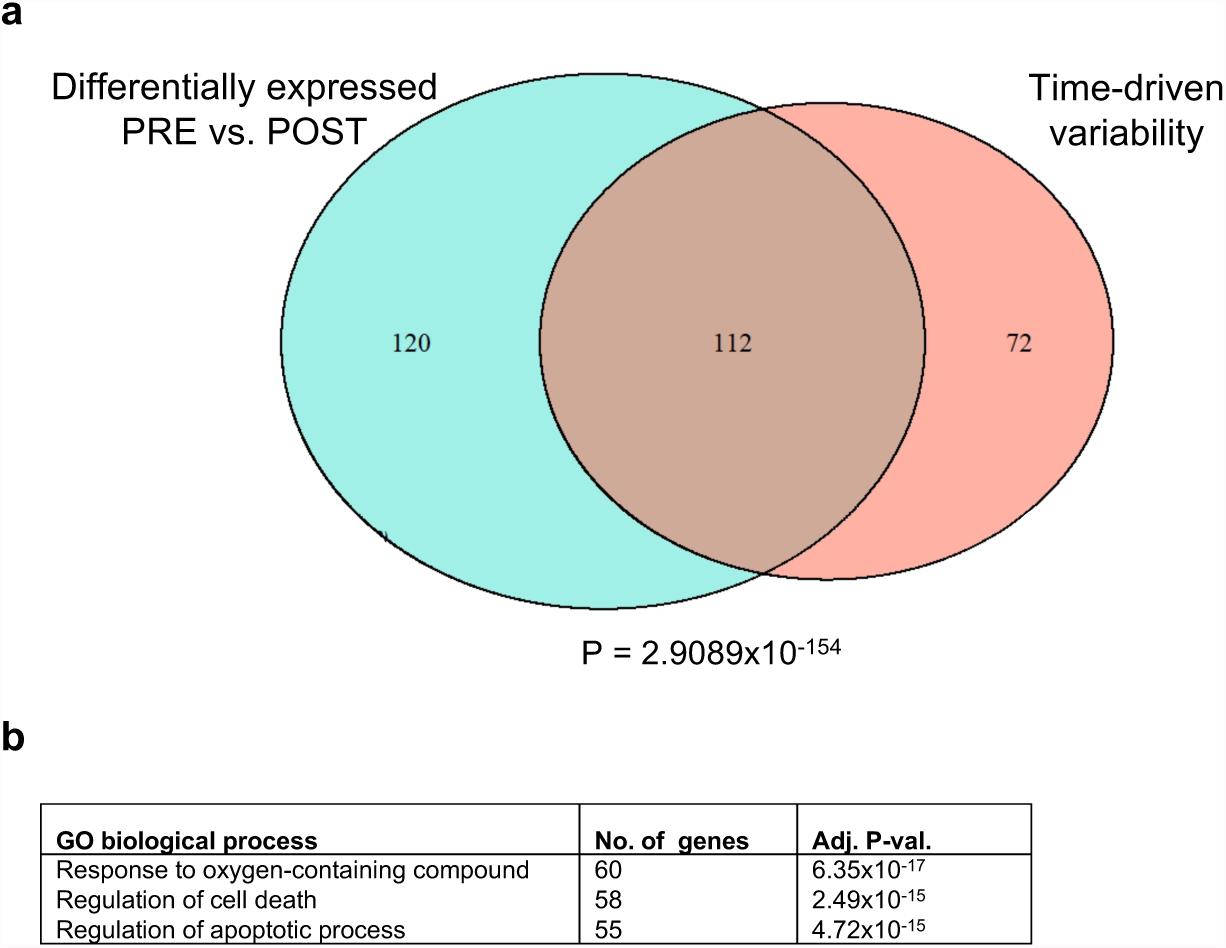
Time-driven variance and early response to ischemia reperfusion. (a)Venn diagram showing the overlap between the genes differentially expressed in POST (compared to PRE) and genes displaying a time-dependent variance (as shown in Fig. 1d). (b) List of most significant GO terms (biological process), according to the gene enrichment analysis including genes displaying a time-driven variance (as shown in Fig. 1d).

**Suppl. Fig. 3.**
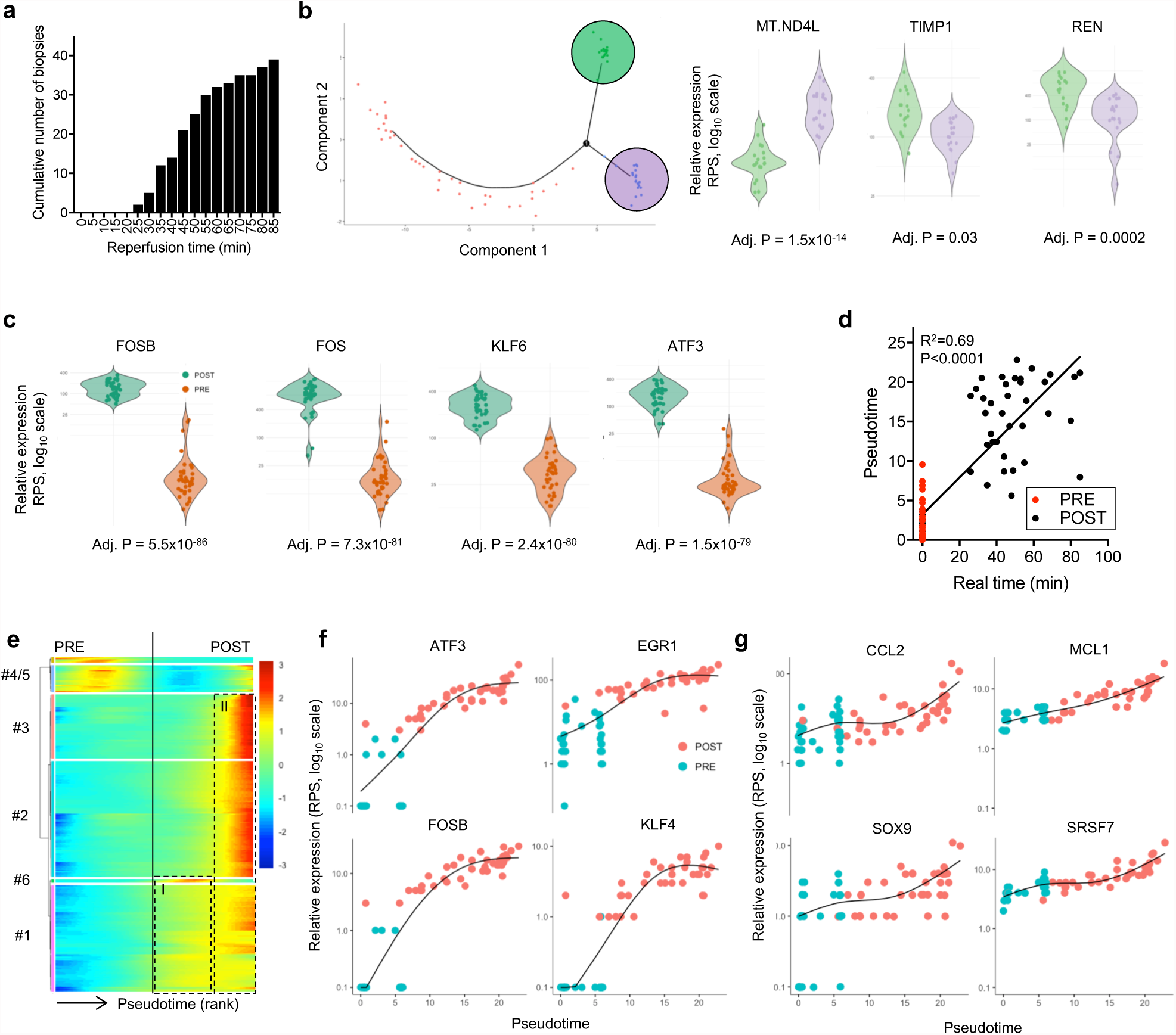
Early response to ischemia reperfusion injury. (a) Histograms indicating the cumulative reperfusion time distribution according to clinical records. (b) Violin plots showing representative examples of genes differentially expressed in the two groups of samples identified by the Monocle algorithm in the PRE samples. The relative expression in RPS is shown for the two groups identified by colors as indicated on the plot on the left (s. Fig. 2a). Adjusted P values are reported (BenjaminiHochberg). (c) Violin plots of the top 4 up-regulated genes in POST compared to PRE samples. Adjusted P values are reported (Benjamini-Hochberg). (d) Dispersion plot showing the correlation between the reperfusion time according to clinical records and the pseudotime calculated by the Monocle algorithm. (e) Cluster analysis of all genes differentially expressed along the pseudotime presented in Fig. 2a: samples are aligned from left to right according to the order shown in Fig. 2a. Genes are vertically aligned and classified in clusters as indicated on the left. The colors indicate the relative expression of the genes. I and II indicate the 2 waves of genes up-regulated after reperfusion (vertical line). The complete list of genes is presented in Suppl. Table 1 (f-g) Representative examples of genes upregulated in the first (f) and in the second (g) wave of transcriptional regulation after reperfusion.

**Suppl. fig. 4.**
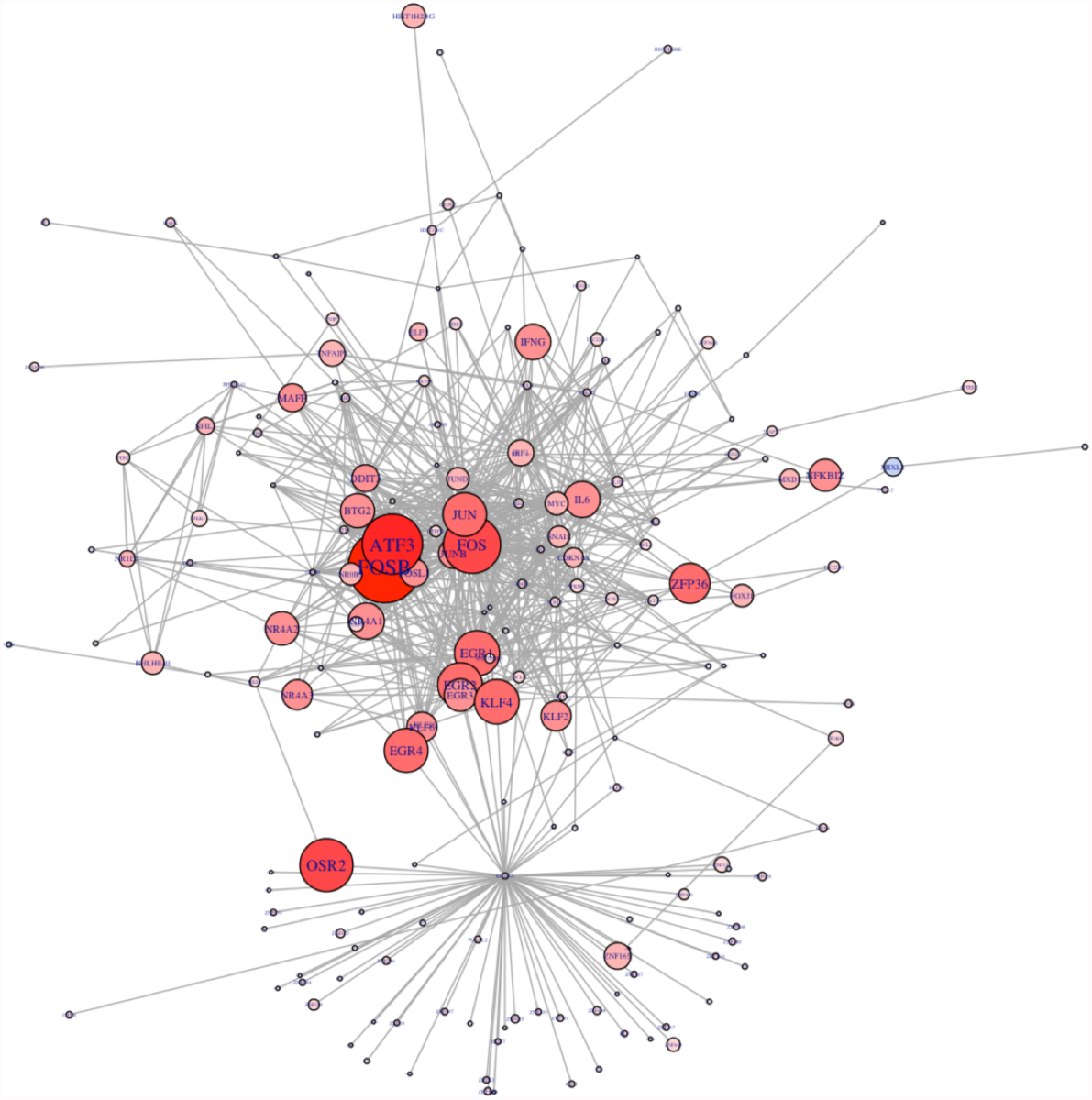
Network analysis. Kamada Kawai plot (igraph network visualization) of the transcription factor centered gene network analysis based on differentially expressed genes between PRE and POST. The size of the vertices correlates with the absolute value of log fold change. Red indicates upregulated in POST. The top 200 genes are shown.

**Suppl. fig. 5.**
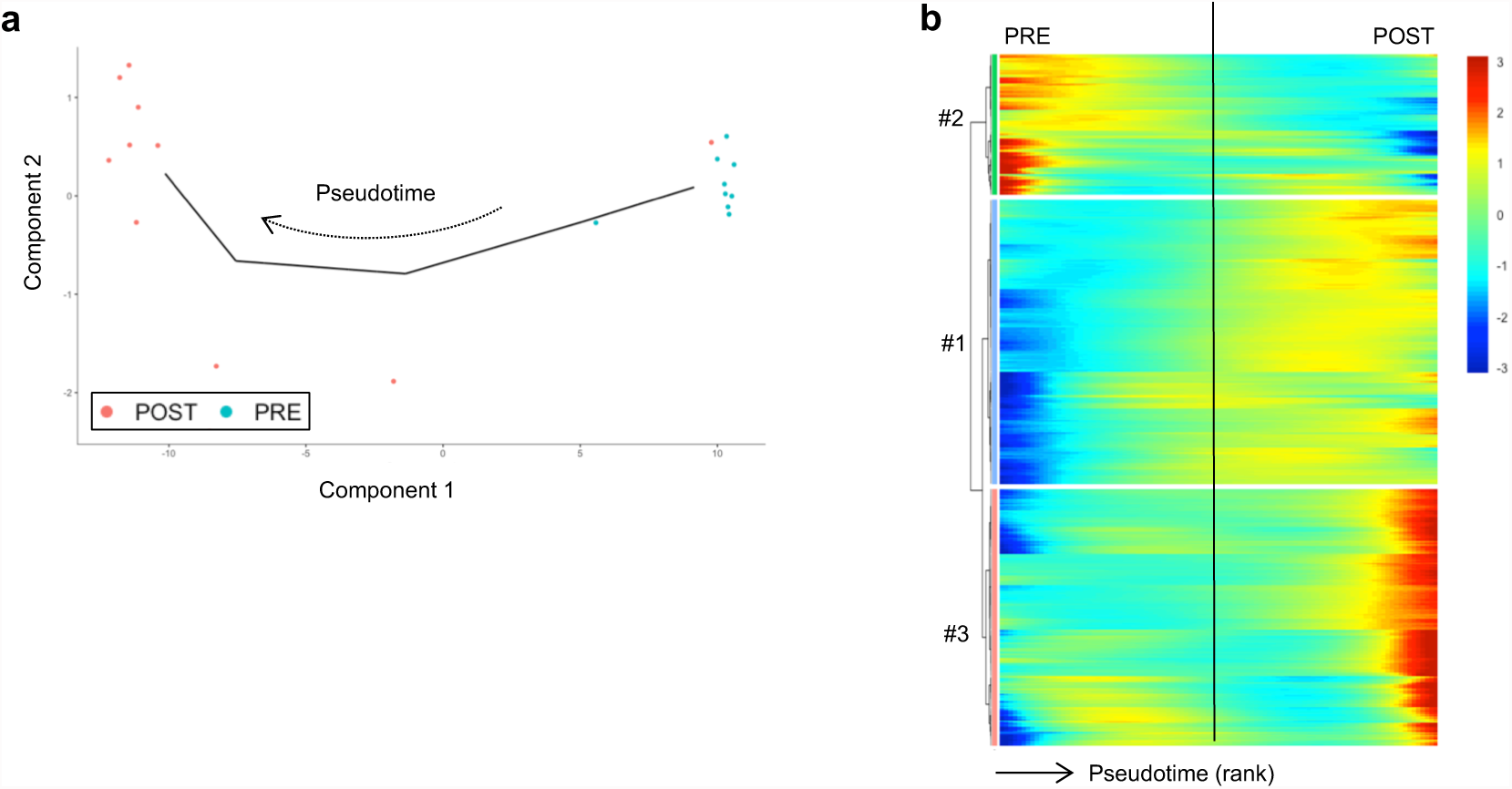
Early response to ischemia reperfusion in the human liver. (a) Sample state ordering in the reduced dimensional space of PRE and POST liver samples, as determined by the Monocle algorithm. (b) Cluster analysis of all genes differentially expressed along the pseudotime presented in (a). Samples are aligned from left to right according to the order shown in (a). Genes are vertically aligned and classified in clusters as indicated on the left (the list of genes is presented in the supplementary materials). The colors indicate the relative expression of the genes.

**Suppl. fig. 6.**
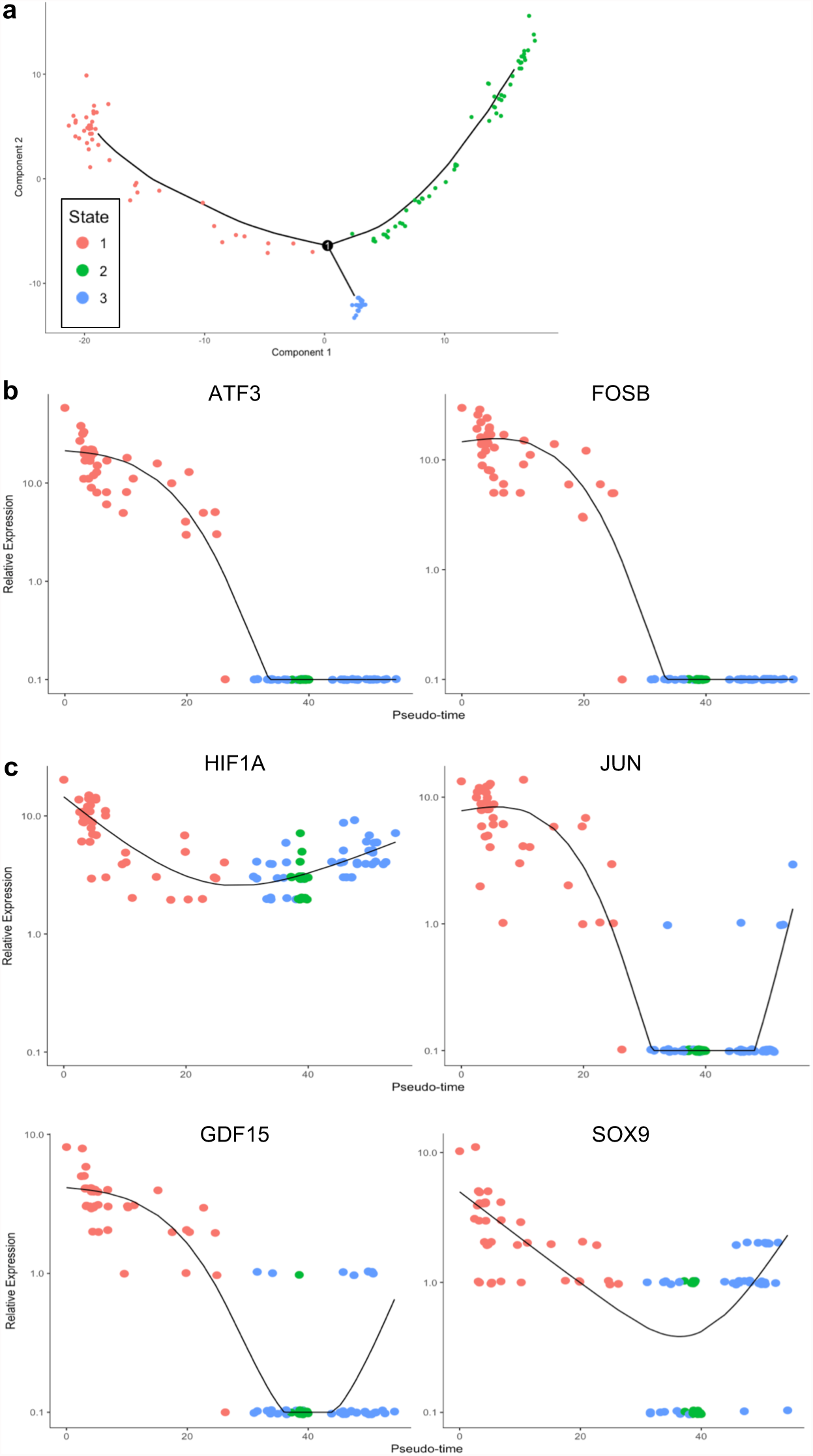
Late regulation of genes involved in the early response to ischemia/reperfusion injury. (a) Pseudotime analysis as reported in Fig. 3e, indicating the 3 states identified by the Monocle algorithm (shown here as a reference for the interpretation of the following plots). (b-c) Representative examples of the relative expression of early response genes along the pseudotime. All early-response genes were strongly up-regulated in the POST samples;some genes were consistently low at 3 and 12 months (b), whereas another set of genes was up-regulated in the late phases of the transition to fibrosis (c).

